# Characterization of siderophore producing arsenic-resistant *Staphylococcus sp*. strain TA6 isolated from contaminated groundwater of Jorhat, Assam and its possible role in arsenic geocycle

**DOI:** 10.1101/231241

**Authors:** Saurav Das, Madhumita Barooah

## Abstract

The presence of arsenic in sediments, carbonaceous rocks are geogenic, while its entry into the aquifers is mediated by several factors including microorganisms. It is well known that the microorganisms play a crucial role in the biogeochemical cycle of different elements. However, the precise role of bacteria in regulating the concentration of arsenic in Brahmaputra valley has not been investigated in detail. In this paper, we report the isolation of arsenic resistant bacterium TA6 with active arsenate reduction efficiency. The isolate was able to grow in arsenate concentration (250 mM) and arsenite (30 mM). Along with resistance to inorganic arsenic, it showed cross-tolerance to other heavy metals like Hg^+2^, Cd^+2^, Co^+2^, Ni^+2^, Cr^+2^. The bacterium also had a high siderophore activity (78.7 ± 0.004 μmol), which is positively correlated with the resistance aptitude. The biochemical test showed the TA6, a gram-positive bacterium which can hydrolyze starch and casein, produce catalase enzyme and utilizes citrate as a metabolic trait. Molecular and chemotaxonomic identification of TA6 based on 16S rRNA and FMAE analysis showed similarity with members of *Staphylococcus* genus with significant difference in sequence similarity and fatty acid composition. Based on 16S rRNA and FAME analysis it was identified as *Staphylococcus sp.* TA6. Rate of biotransformation showed bacterium could reduce ~88.2% of initial 2mM As(V) into As(III). The characterization of arsenate reductase enzyme with NADPH coupled assay showed the highest activity at pH 5.5 and temperature 50°C.

## 1.0 INTRODUCTION

Brahmaputra river basin is long known for its high geogenic arsenic (As) concentration. However, during the recent times, the problem has much aggravated as 23 out of the 32 districts in Assam have been reported to be affected by high Arsenic concentration [1,2]. Titabor subdivision of Jorhat district is considered as one of the most As affected areas of Assam. The concentration of arsenic in this region ranges between 194-657 μg/l, which is far above the permissible standard of BSI (50 ng/l) and WHO (10 ng/l) [3]. Arsenic is a metalloid widely distributed in the earth’s crust and its concentration can exist from traces to up to hundreds of mg/kg or mg/l in both soil and in water. In groundwater, the element is predominantly found in two states *viz.*, arsenate (AsV) and arsenite (AsIII). Arsenate is dominant in the oxic environment and gets strongly absorbed by chemicals like ferric-oxyhydroxide, ferrihydrite, apatite, and alumina. The arsenite form is dominant in an anoxic environment and is more mobile and toxic than arsenate [4]. The geochemical cycling of arsenic is composite in nature; in fact, in addition to various physical and chemical factors, it also involves biological agents. Microorganisms play a critical role in mobilization and speciation of arsenical compounds in aquifer systems [5]. Microorganisms in the course of evolution have developed the necessary genetic makeup which confers them with resistance to high concentration of arsenic as well other toxic metalloids [6]. Microorganism arbitrated mechanisms of arsenic mobilization are still poorly understood and therefore, needs further studies to reveal their role in sediment-bound arsenic mobilization. They can either reduce, oxidize or can methylate the element to other organic compounds to generate energy [7]. Arsenate reducing bacteria are able to reduce As (V) to As (III) and use the same as an electron acceptor in a respiratory pathway or efflux the same as a mean of resistance mechanisms [8]. Arsenic resistant bacteria are frequently detected with siderophore activity. Siderophore are high-affinity iron chelating compounds produced and secreted by few microorganisms to forage the environmental iron from inorganic phase by formation of soluble Fe^3+^ complex, which can be taken up by active transport mechanisms [9]. The Fe sequestering ability of bacteria through siderophore production confers them with an added advantages over the non-producers in arsenic resistance. The previous study has shown that the rate of arsenic uptake and reduction efficiency of a bacteria significantly varies with varied siderophore concentration [10].

In this paper, we report the isolation of a gram-positive bacterium TA6, which is resistant to both arsenate and arsenite. Based on its morphological, molecular and chemotaxonomic characterization the isolate was identified as a species of the genus *Staphylococcus* that had facilitated arsenate reduction along with siderophore activity.

## 2.0 MATERIALS AND METHODS

### 2.1 Sample Collection

Contaminated groundwater samples were collected from Tanti-Gaon (GPS: 26.58.101, 94.16.391) (**Fig. 1**), a village in Titabor subdivision of Jorhat district, Assam, India. The concentration of arsenic in the water samples was measured by atomic absorption spectrophotometer (AAS) following the standard protocol as described by Behari and Prakash [11].

**Figure 1:**
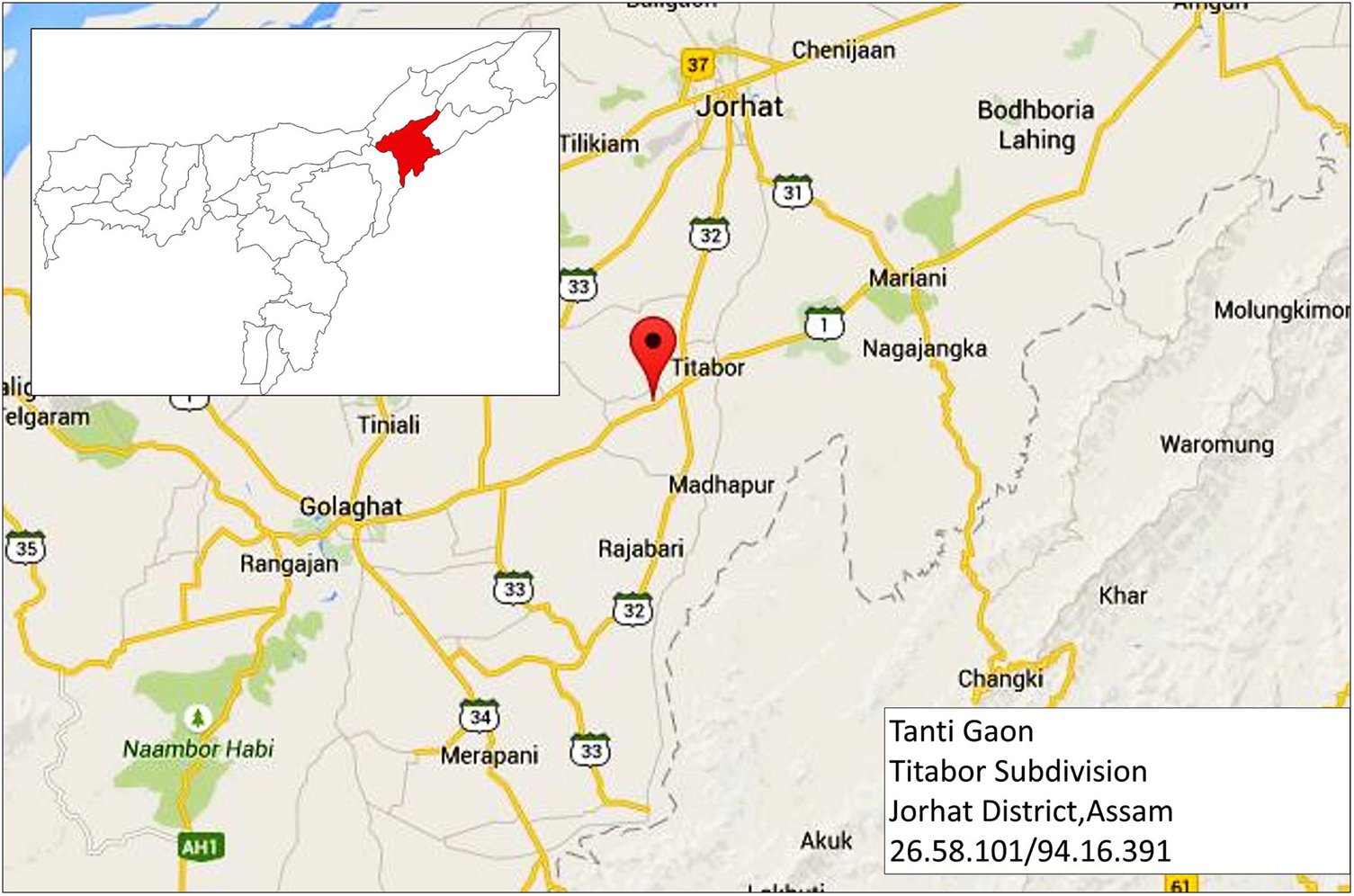
Site map of the study area.

### 2.2 Isolation of arsenic-tolerant bacteria

The collected sample was enriched in LB broth, subjected to serial dilution and cultured in arsenate amended LB agar plates (10mM Arsenate/1 mM of Arsenite) and incubated at 30°C for 48 hrs. Individual colonies were picked up based on the morphological identities and sub-cultured to obtain the pure isolates.

### 2.3 Minimum Inhibitory Concentration Test

The minimum inhibitory concentration (MIC) of arsenate [As (V)] and arsenite [As (III)] was evaluated to determine the resistance capacity of the isolated bacteria. The bacterial isolates were cultured in freshly prepared LB broth at 30°C for 48 hours and then 100 μl of the freshly cultured bacterial suspension was inoculated in minimal salt media (MSM) supplemented with different concentration of arsenite (0.5-30 mM) added as sodium meta-arsenite (m-Na-AsO_2_) and arsenate (10-300 mM) added as disodium hydrogen arsenate (Na_2_HAsO_4_.7H_2_O) and incubated for 72 hours at 30^0^ C and 142 rpm. The microbial growth was recorded with a UV-Visible spectrophotometer at 600nm.

### 2.4 Growth of the bacterial isolate in the presence and the absence of arsenite/arsenate

Among all the isolates, TA6 showed the highest MIC and as such, was taken for studying the growth kinetics in presence and absence of arsenite and arsenate. The isolate was cultured in Luria-Bertani broth containing arsenate in a concentration of 1mM to 30mM and arsenite from 0.5 mM to 10 mM respectively. The growth of the isolate was monitored through measurement of the optical density (OD) with a spectrophotometer (Thermo-Scientific, India) at 600 nm (OD600) at a specified interval of time (4hr, 8hr, 12hr, 24hr, 48hr, and 72hr).

### 2.5 Biotransformation assay

The ability of the bacteria to reduce As (V) or to oxidize As (III) was evaluated using the silver nitrate (AgNO_3_) method as described by Simeonova *et al.*,[12]. Freshly cultured bacterium grown in minimal salt medium with 5mM glucose was subcultured on two different LB agar plates supplemented with 2mM of Sodium Meta-Arsenite and Sodium Arsenate respectively and incubated for 48 hours at 30°C. The streaked plates were then flooded with 0.1M Silver Nitrate (AgNO_3_) solution. Formation of light yellow color will indicate the precipitation of silver ortho-arsenite (Ag3AsO3) and light brown-red color for precipitation of silver-ortho-arsenate (Ag3AsO4).

Quantitative assay of arsenate reduction was analyzed by culturing the bacteria in arsenic amended LB broth (2mM of Arsenate). In a time interval of 6, 12, 24, 48, 72 hours the bacterial cells were collected by centrifugation and arsenite content of the supernatant was determined by AAS following standard protocols as described by Aggett and Aspell [13].

### 2.6 Arsenate reductase enzyme assay

The enzyme assay was done using NADPH coupled assay as described by Gladysheva *et al.*, [14]. Cell-free crude extracts of *Escherichia sp.* SD23 was used as positive control. Effect of pH and temperature on enzyme activity was also measured using this method.

### 2.7 Cross tolerance

The isolate was tested for its cross-tolerance efficiency with other heavy metals like Hg^+2^ added as HgCl_2_, Cd^+2^ added as CdCl_2_, Co^+2^ added as CoCl_2_, Ni^+2^ added as NiCl_2_ and Al^+3^ added as AlCl_3_ in a concentration ranging from 0.5 to 10mM in MSM broth culture and OD (OD600) was recorded after 48 hours to evaluate the bacterial growth.

### 2.8 Biochemical Tests and Carbon Source Utilization

Biochemical tests for starch hydrolysis, catalase, oxidase, casein production, nitrate reduction, urease, malate, citrate, indole, and motility were done according to the standard protocol described by Krieg [15]. Carbon source utilization was tested using BioMerieux 50CHB/E strips (BioMerieux, USA)

### 2.9 Identification based on 16S rRNA and Phylogeny

Genomic DNA was extracted from approximately 100 mg of the cell as per standard phenol-chloroform method. The 1500bp region of the 16S rRNA gene was amplified from the extracted genomic DNA using the universal primer 5′ TACGGYTACCTTGTTACGACTT 3′ (1492R), 5′ AGAGTTTGATCMTGGCTCAG 3′ (27F). The amplification was carried out in a reaction with a final volume of 25 μl containing 1.5 μl of template DNA, 1 μl (20pM) of the forward primer, 1 μl (20 pM) of the reverse primer, 2.5 μl (2.5 mM of each) dNTP mix, 2.5 μl of 10x PCR buffer, 1 μl (1U) of Taq DNA polymerase. A negative control (PCR mix without DNA) was included in all PCR experiments. The PCR reaction conditions were set for 94°C for 3min, followed by 30 cycles of denaturation at 94 °C for 30s, annealing at 58°C for 1min and extension at 72°C for 2 min, before a final extension at 72 °C for 7min. The PCR products were sequenced and the forward and reverse sequences obtained were assembled using the Codon-Code Aligner software (version: 5.1). Nucleotide sequence identities were determined using the BLAST tool from the National Center for Biotechnology Information (NCBI) and Similarity index value from EzTaxon Server. The partial sequence data for the 16SrRNA genes have been submitted to GeneBank for further references. Phylogenetic relationship inferred with neighbor-joining (NJ) tree [16]. Sequence divergence among the strains was quantified using Jukes-Cantor distance model [17]. A total of 1000 bootstrap replication were calculated for evaluation of the tree topology.

### 2.10 FAME analysis

The fatty acid methyl ester (FAME) profile was analyzed using Sherlock-Midi system and compared with few reference strains of *Staphylococcus* genus for taxonomical validation.

### 2.11 Siderophore production

Production of siderophore was studied using Chrome Azurol S (CAS) agar media as described by Schwyn and Neilands [18]. CAS agar was prepared from four solutions which were sterilized separately before mixing. The **solution I**: Blue dye was prepared by mixing 10 ml of 1 mM FeCl_3_, 6H_2_O in 10mM HCl then with 50 ml of an aqueous solution of 2mM CAS. The resulting dark purple mixture was added slowly with constant stirring to 40 ml of an aqueous solution of 5mM Hex-Decyl Tri-Methyl Ammonium [HDTMA]. The dark blue solution was produced which was autoclaved and then cooled to 50^0^ C. All reagents in the indicator solution were freshly prepared for each batch of CAS agar. **Solution II**: CAS agar was prepared by dissolving 30.24 g of Piperazine-N, N’-bis ethane sulfonic acid (PIPES) in 750 ml of a salt solution containing 0.3 gKH_2_PO_4_, 0.5g NaCI, and 1.0g NH_4_Cl. The pH adjusted to 6.8 with 50 % KOH, and water was added to bring the volume to 800 ml and autoclaved after adding 15g of agar, and then cooled to 50^0^ C. **Solution III**: Mix Solution containing the followings: 2g glucose, 2g mannitol, 493 mg MgSO_4_,7H_2_O, l mg CaCl_2_, 1.17 mg MnSO_4_H_2_O, 1.4 mg H_3_BO_3_, 0.04 mg CuSO_4_.5H_2_O, 1.2 mg ZnSO_4_,7H_2_O, and 1.0mg Na_2_MoO_4_.2H_2_O, was autoclaved, cooled to 50°C then added to the buffer solution along with 30 ml filter-sterilized 10 % (w/v) casamino acid (Solution IV). The indicator solution was added last, with sufficient stirring to mix the ingredients without forming bubbles. CAS agar plates were inoculated with bacterial isolate and incubated at 30^0^ C for 7 days. Colonies showing orange hollow zone following incubation were recognized as siderophore positive [19].

### 2.12 Siderophore quantification

The method of Alexander *et al.* [20] was used to measure siderophore production in vitro. The bacterial cells were grown at 30°C for 24 h in 50 ml of Chrome Azurol S (CAS) medium with 5 mM MES (2-(N-morpholino ethanesulfonic acid)– KOH buffer at pH 6.8. After the culture growth attains exponential phase at OD-600, the cells were pelleted by centrifugation at 10,000g for 10 min and the supernatant was filtered through 0.25 μm filter. Siderophore concentration in the filtrate was measured by mixing 500 μl of modified CAS assay solution with 500 μl filtrates. The standard solution of deferoxamine mesylate was used for siderophore quantification. The sterile CAS-MES-KOH solution was used as a reference solution, which did not contain siderophores. A standard curve was prepared by analyzing the absorbance (630 nm) of the reference solution (A/Aref) as a function of the siderophore concentration.

### 2.13 Resistance to Arsenic in comparison to siderophore mutant

The role of siderophore in arsenic tolerance was determined following the protocol described by Ghosh *et al.,* [10] using one siderophore mutant (non-producer) *Pseudomonas putida* (Lp10L02M) and one control *Acinetobacter guillourie* (S02Ar2) with low siderophore production ability (10.8μmol). Arsenic tolerance of the isolate was measured as a percentage of growth rate and As(V) reduction at 5 and 10 mM of As(V) modified LB medium incubated at 30°C for 24 h shaking at 142 rpm and compared with the TA6. Growth was measured as OD at 600nm on UV–Vis spectrophotometer. All the data were taken in triplicates.

## 3.0 RESULTS

### 3.1 Groundwater Sample

The contaminated groundwater samples collected from Titabor subdivision had pH 6.2 –7.3 and arsenic concentration of 50 - 356 μg/l.

### 3.2 Isolation of arsenic-resistant bacteria and MIC

The enriched groundwater sample was inoculated in arsenate amended LB medium and morphologically different bacterial colonies were picked up and tested for minimum inhibitory concentration of arsenate and arsenite. Among the isolates, TA6 showed highest MIC and was able to grow in medium with 250mM of arsenate and 30mM of arsenite.

### 3.3 Bacterial growth in presence of arsenate and arsenite

Growth curve analysis showed the effect of arsenate and arsenite in the bacterial growth pattern. The isolate TA6 was cultured in fresh LB broth with a concentration of arsenate varying from 1mM – 30mM and arsenite from 0.5mM – 10mM respectively. Bacterial growth was not much affected in the presence of arsenate as compared with control. However, the presence of arsenite in the medium greatly affected the rate of growth. In the presence of arsenate, TA6 started doubling at the lowest time of 4hrs but in the presence of arsenite, it took approximately 24hrs to start multiplying. At the highest concentration of arsenate (~30mM) taken for the test and at 72 hours of incubation time, OD was measured as 1.474 ± 0.067, which is statistically at par with the OD 1.962 ± 0.058 for control, at the same time of incubation. While, at 72hrs of incubation in the presence of 10mM of arsenite growth was reduced when compared to the control. For control, OD was recorded as 1.962 ± 0.058, whereas in the 10mM of arsenite, the growth was recorded as OD 0.1036 ± 0.043. At lowest concentration of arsenite 0.5mM, the bacterial cell (TA6) approximately took 8 ± 2 hours of incubation to multiply (**Fig. 2**).

**Figure 2:**
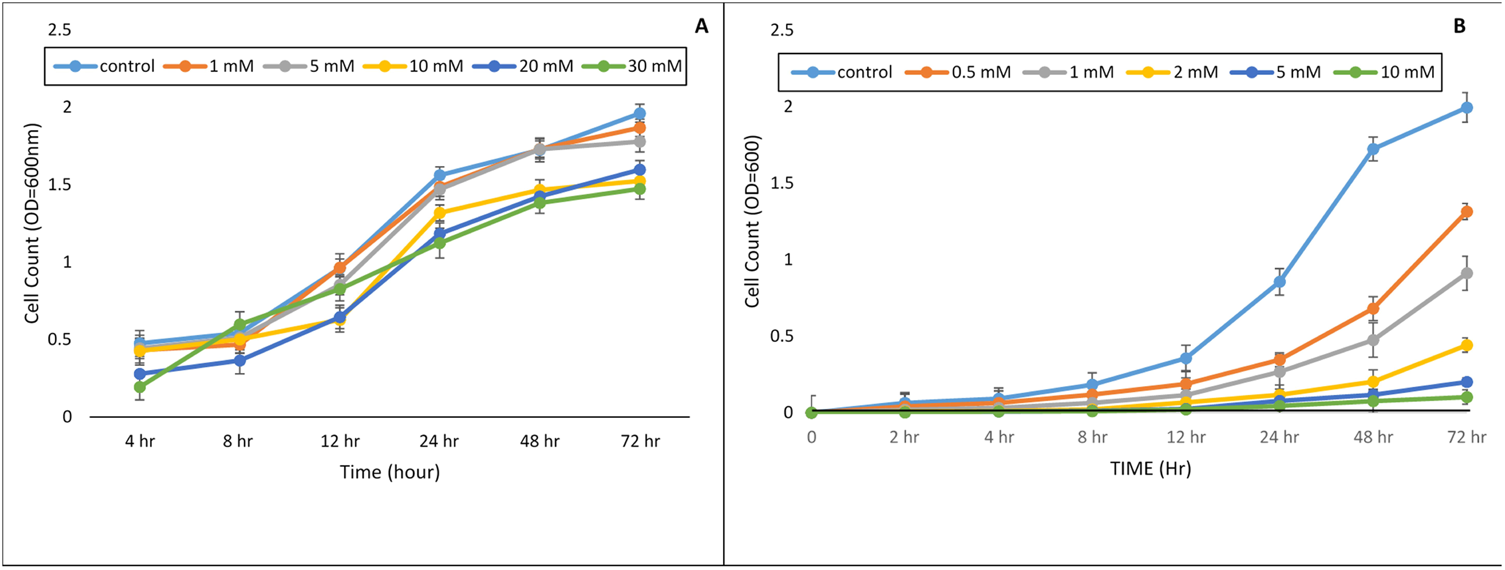
Effect of arsenic on bacterial growth rate (A) Arsenate (B) Arsenite.

### 3.4 Cross tolerance

Other heavy metal tolerance test also showed the resistive capacity of the isolate to various heavy metals like Hg+2, Cd^+2^, Co^+2^, Ni^+2^, Cr^+2^. MIC was found as 0.5mM, 0.8mM, 1.0mM, 4mM, and 6mM respectively.

### 3.5 Biotransformation assay

TA6 was found to be an arsenate reducer. Reduction of arsenate in the petri dish formed a yellow precipitation of silver ortho-arsenite (Ag_3_AsO_3_) which indicates the presence of arsenite. In the quantitative assay, it was also found that with a gradual increase in time and with the increased bacterial cell count, the concentration of As(V) gradually decreased with increased concentration of AS(III). In a duration of 72 hours, nearly 88.2% of the initial 2mM As(V) is reduced to As(III) (**Fig. 3**).

**Figure 3:**
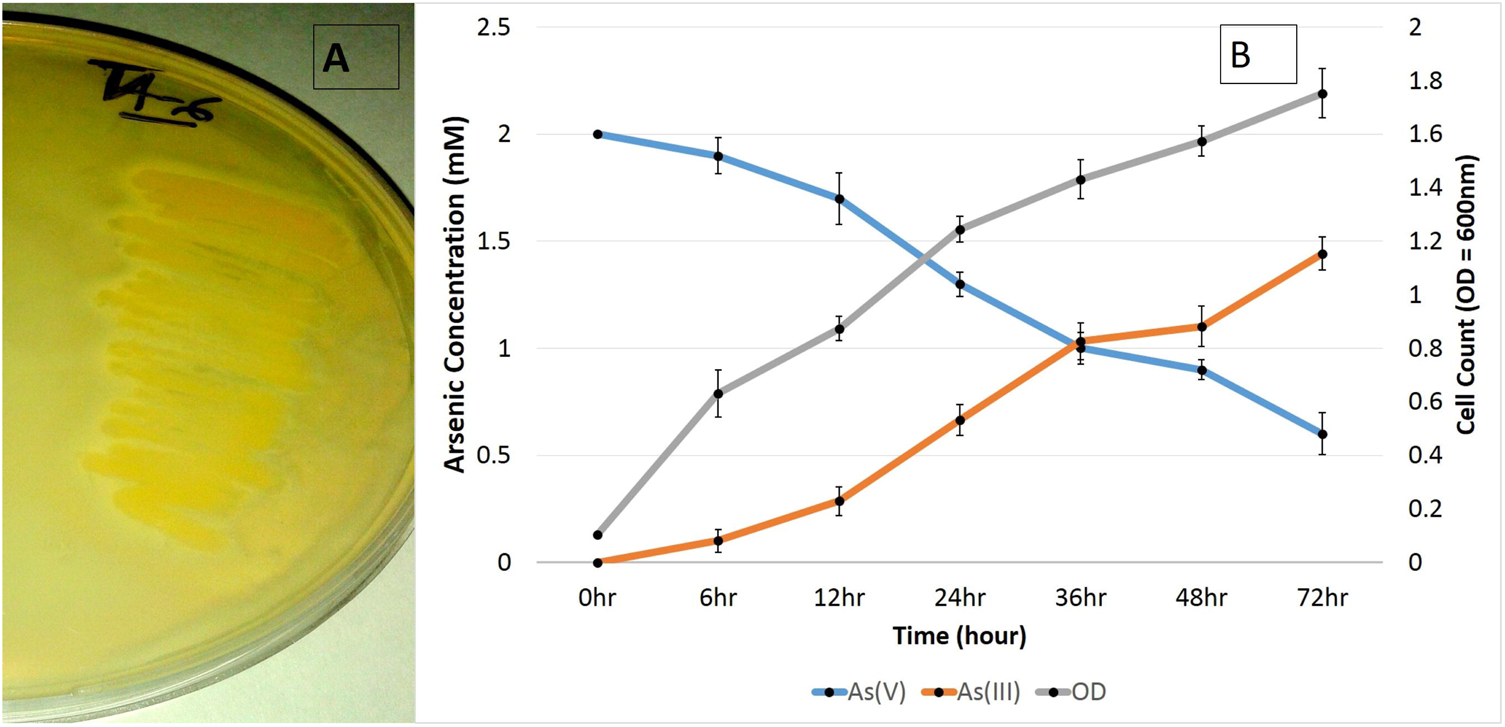
**(A)** Bioconversion of arsenate [As (V)] to arsenite [As (III)]. (**B**) Rate of Biotransformation over an incubation period of 72 hours.

### 3.6 Arsenate reductase enzyme activity

Arsenate reductase activity was measured using NADPH coupled oxidation method. A Km of 0.44 mM arsenate and Vmax of 6395 Umol/min were measured (**Fig. 4**). There was no change in activity for 500μM and 1mM of arsenate. Temperature and pH are some critical factors for enzyme activity. Temperature-dependent activity assay revealed that 50°C was the optimal temperature for highest enzymatic activity and in pH-dependent activity assay, pH 5.5 was measured as optimal for highest enzymatic activity (**Fig. 5**). Graphical representation of both the data formed a characteristic bellshaped curved, where initial increased pH and temperature raised the activity till it reaches the optimal point of maximum activity and then the activity was found to gradually cease after the respective optimal value of pH 5.5 and temperature 50^0^ C.

**Figure 4:**
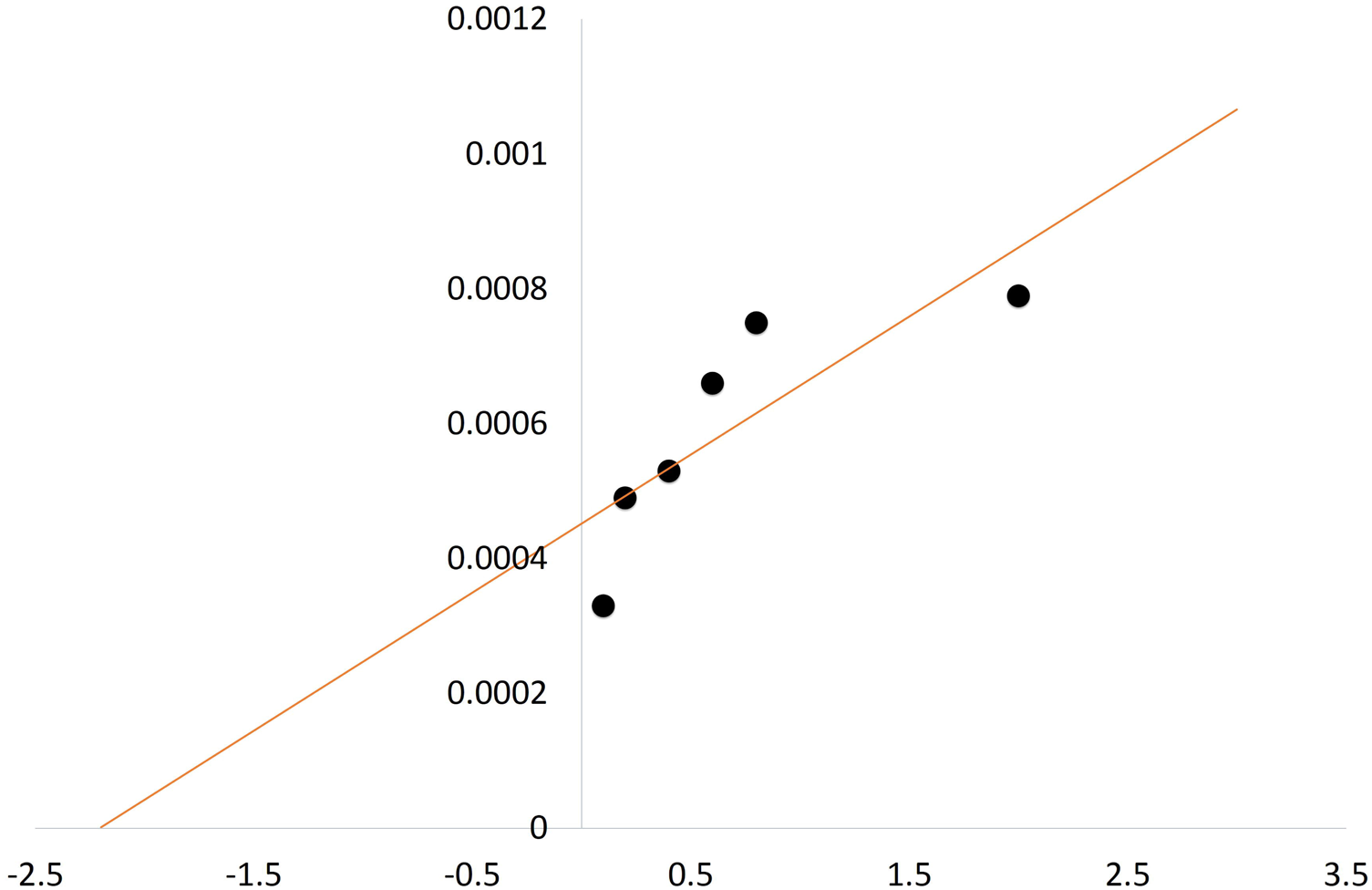
Kinetic Profile of enzyme activity (Lineweaver Burk Plot).

**Figure 5:**
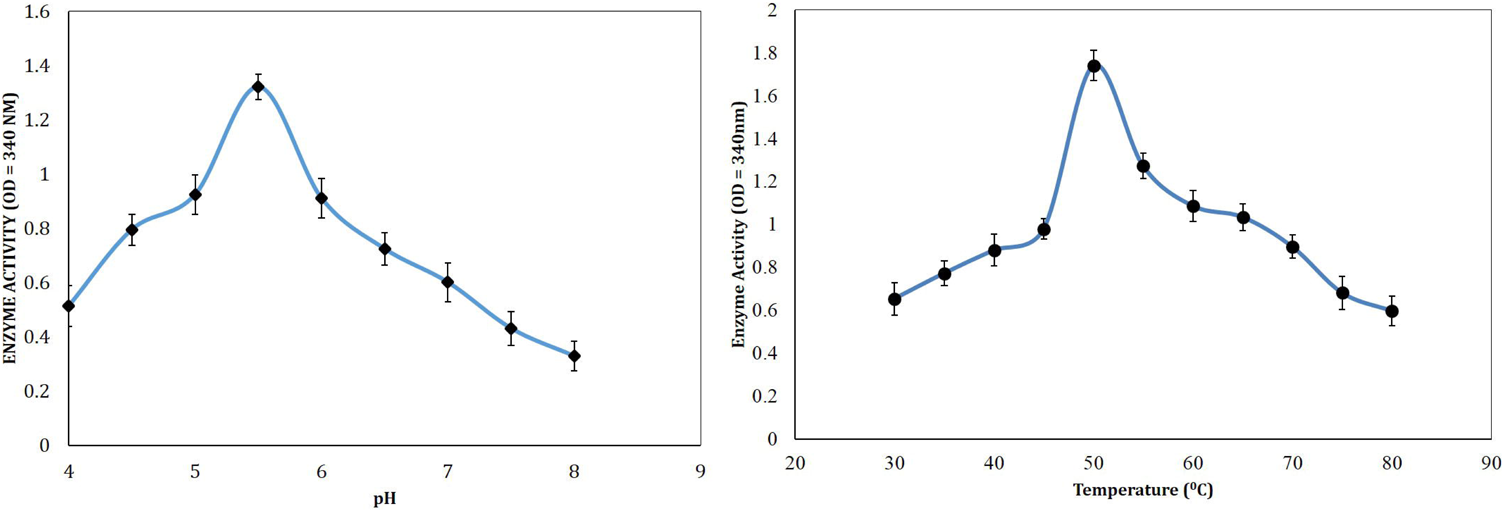
Arsenate Reductase enzyme activity (**A**) At different pH (pH 5.5 was found to be optimal for maximum activity) (**B**) At different Temperature (50 °C was found as the optimal temperature for maximum activity).

### 3.7 Biochemical test

The bacterium (TA6) was a gram-positive, non-motile, coccus shaped bacterium. It was able to hydrolyze starch, casein and utilize citrate, reduce catalase and showed high siderophore activity (78.7 ± 0.004 μmo1) but tested negative for oxidase, nitrate, urease and indole. Carbohydrate utilization test with 50CHB/E showed it could actively utilize Glycerol, D-Glucose, D-Fructose, Maltose, Lactose, Sucrose, Trehalose, Melezitose, Starch, and D-Turanose.

### 3.8 Chemotaxonomic and Molecular Identification with Phylogeny

The 16S rRNA sequence similarity search identified the isolate as one of the species of the genus *Staphylococcus* having 98% pairwise similarity with *Staphylococcus saprophyticus subsp. Bovis* MM19 and *Staphylococcus saprophyticus* strain OUCMDZ4189. Fatty acid methyl ester profile showed most of the fatty acids are branched chain like anteiso-C15, anteiso-C17 and iso-C15, when compared with the fatty acid profile of *S. xylosus, S. cohnii and S. saprophyticus* showed considerable differences of C17:0, iso C17:0, iso C18:0 (**Table 1**). Therefore, based on both molecular and chemotaxonomic data the bacterium was identified as *Staphylococcus sp.* and the sequence was submitted under the GeneBank accession: KF134542.1 for further references. Phylogenetic analysis showed significant evolutionary difference among the other member of the *Staphylococcus* genus but with similar lineage of origin (**Fig. 6**). Evolutionary distance computed with Jack Cantor model and 1000 bootstrap value showed TA6 is 68% similar on evolutionary lineage with *Staphylococcus saprophyticus subsp. Bovis* MM19.

**Table 1:** Cellular Fatty acid Profile of iso1ateTA6 (1) and *S. xylosus* (2) *S. cohnii* (3) and *S. saprophyticus* (4).

| 433 | **Straight chain fatty acids** | 1 | 2 | 3 | 4 |
| --- | --- | --- | --- | --- | --- |
| 434 | C15:0 | -- | -- | -- | -- |
| 435 | C16:0 | 1.61 | 3.44 | 3.21 | 3.57 |
| 436 | C17:0 | 0.07 | -- | -- | -- |
| 437 | **Branched chain fatty acids** |  |  |  |  |
| 438 | iso C14:0 | 0.96 | 2.40 | 2.48 | 1.88 |
| 439 | iso C15:0 | 14.04 | 23.73 | 15.01 | 21.09 |
| 440 | anteiso- C15:0 | 50.58 | 39.24 | 43.28 | 41.55 |
| 441 | iso C16:0 | 2.06 | 0.97 | 3.14 | 1.94 |
| 442 | iso C17:0 | 6.80 | -- | -- | -- |
| 443 | anteiso - C17:0 | 13.13 | 4.48 | 10.18 | 4.66 |
| 444 | iso C18:0 | 0.47 | -- | 0.62 | -- |

**Figure 6:**
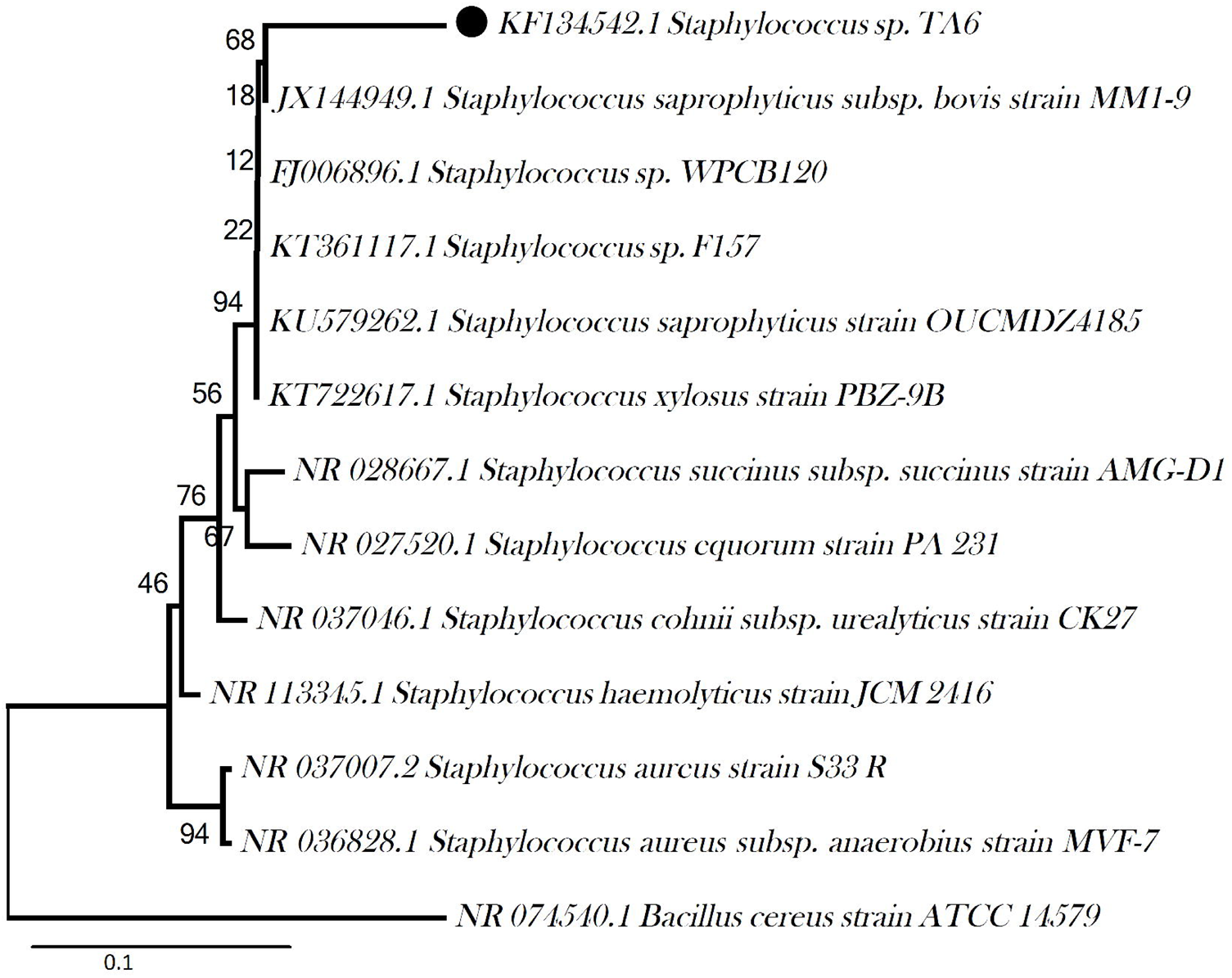
The evolutionary history was inferred using the Neighbor-Joining method. The optimal tree with the sum of branch length = 0.90052 (~ 0.1) is shown. The percentage of replicate trees in which the associated taxa clustered together in the bootstrap test (1000 replicates) are shown next to the branches. The tree is drawn to scale, with branch lengths in the same units as those of the evolutionary distances used to infer the phylogenetic tree. The evolutionary distances were computed using the Jukes-Cantor method and are in the units of the number of base substitutions per site. All positions containing gaps and missing data were eliminated.

### 3.9 Siderophore associated arsenate reduction

Microorganisms are the primary chelator of iron which dissociates Fe^+3^ ions with their siderophore activity. Siderophore associated arsenic resistance assay revealed that bacteria with high siderophore TA6 (78.7 ± 0.004 μmol) was significantly a strong As(V) reducer than the mutant strain Lp10L02M (non-producer). The growth of TA6 was also found reflective in comparison to the control and the mutant implying the added resistance ability of the strain to arsenate. In 5mM and 10mM arsenate broth, the TA6 showed higher growth as compared to the control strain S02Ar2. However, the mutant strain (Lp10L02M) had slower growth rate as compared to control and showed lesser reduction efficiency (Fig. 7).

**Figure 7:**
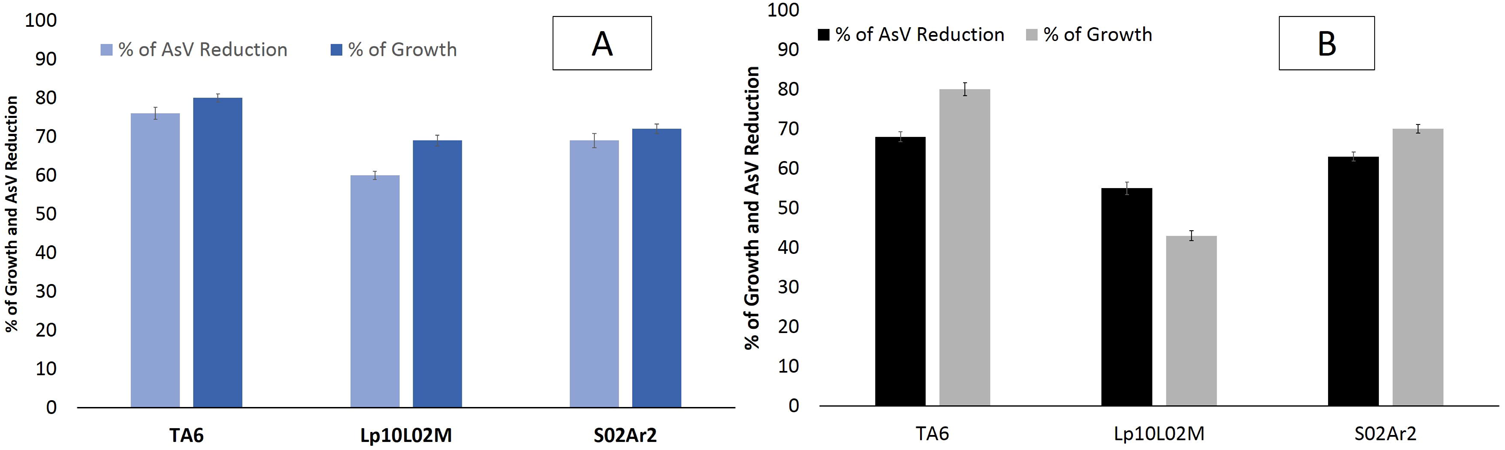
Effect of siderophore on growth percentage and arsenate reduction in the modified LB medium containing (A) 5mM arsenate (B) 10mM arsenate.

## 4.0 Discussion

Increased arsenic concentration in groundwater is a matter of serious concern due to its carcinogenic effects on human health. The Brahmaputra river basin is considered as one of the severely affected basins [21]. Flood-line areas of the river have been detected with high concentration of arsenic above the standard permissible limit. It has become a major health risk for the people residing within these vicinities as they solely depend on the natural streams and groundwater for potable water. Titabor subdivision of Jorhat district, Assam has been severely affected by an alarming concentration of arsenic (194-657 μg/l)[3]. Although, several studies on arsenic poisoning and geogenic distribution in this region has been documented [22-24], the role of microbes in the geocycle still needs a long way to go. Bacteria play an important role in the biogeochemical cycle of arsenic and they are contemplated as one of the major factors that have a pivotal role in the mobilization of sediment-bound arsenic [25]. Our previous study on microbial diversity of contaminated aquifers proves the same [26]. They can change the bioavailability, solubility and mobility of arsenical compounds by inter-conversion through redox reactions and thus, control toxicity level. Bacteria can resist the toxic arsenic using an array of cellular and metabolic mechanisms which basically includes active cellular transportation of the toxic material out of the cellular environment, entrapment by cellular capsules or by precipitation, oxidation-reduction reaction [5,27]. We isolated a bacterium TA6 from groundwater sample containing 356μg/l of arsenic. The bacterium was identified as *Staphylococcus* sp. based on the 16S rDNA sequence analysis and fatty acid methyl ester (FAME) profile. Both 16S rDNA and FAME analysis showed significant differences with the reference strains of *Staphylococcus*. Similarity search with the NCBI nr/nt database and EzTaxon server showed an average of 98% similarity with the different species of *Staphylococcus* genus. Straight chain fatty acids like c16:0, c17:0 and branched chain fatty acids like iso c14:0, iso c17:0, anteiso c15:0 showed considerable difference when compared with the FAME profile of *S. xylosus, S. cohnii* and *S. saprophyticus* (Table. 1). The bacterium was found to tolerate arsenate (MIC: 250 mM), arsenite (MIC: 30 mM). Resistance to arsenite concentration greater than 10 mM and arsenate greater than 100 mM has been considered as very high, whereas resistance to 200 mM As^5+^ and 30 mM As3+ is a hyper-tolerance property [28]. This higher tolerance to inorganic arsenic is may be due to the presence of arsenic resistance operonic genes *(arsR, arsB* and *arsC)* as confirmed by PCR detection method (data not shown). The presence of arsenic resistance genes among the members of *Staphylococcus* genus is well documented [29-31]. The bacterium showed resistance to other heavy metals like Hg^+2,^ Cd^+2^, Co^+2^, Ni^+2^, Cr^+2^ and MIC ranged from >0.5-10.0 mM.

Microorganisms are known for their ability to produce different biogenic chelating agents like siderophore in iron-limiting environment. Siderophore solubilize the ferric iron in the iron-starved environment and transports the Fe^+3^ into the cell [32]. It determines the growth of microorganisms in an environment where iron is the limiting factor [33]. The bacterium was found to produce significantly high amount of siderophore (78.7 ± 0.004 μmol). Above and beyond being a limiting factor for growth in iron-starved environment, siderophore confers an added advantage of higher arsenic resistance as compared to the non-producers [10]. Screening of comparative resistance efficiency TA6 (78.7 ± 0.004 μmo1) with a control strain *Acinetobacter guillourie* S02Ar2 (10.8 ± 0.003 μmo1) and a mutant strain *Pseudomonas putida* (Lp10L02M) showed a significant difference. The bacterium TA6 was able to resist higher concentration of arsenate in comparison to the mutant and control strain. Siderophore assisted resistance to arsenical compounds has been reported earlier [10]. High siderophore concentration confers higher resistance to arsenate and that the reduction efficiency of bacteria is also significantly influenced by varied siderophore concentration.

Biotransformation assay indicated the isolate as one of the members of the arsenate-reducing genera that can actively catalyze the reduction of As(V) to As(III) using an enzyme arsenate reductase encoded by *arsC* gene of the ars operon. Aerobic arsenate reduction is the most distributed detoxification mechanism among the microorganisms and the ars operon has been detected in more than 50 organisms within the domains of bacteria, yeast, and protist. The first recognized arsenate reductase gene was identified in a gram-positive *Staphylococcus* plasmid [27]. Since then there have been several reports of this gene in different bacterial species *viz. Staphylococcus sp., Thermus thermophiles* [34] *Bacillus sp., Shewanella sp.*, [27]. Analysis of NADPH coupled assay showed that the enzyme is slightly acidic in nature with optimal pH 5.5 and optimal temperature 50°C for its highest activity. Michaelis Menten kinetic constant, km was found to be 0.44 mM arsenate and V_max_ of 6395 Umol/min. A similar kinetics of this enzyme was reported from *Chrysiogenes arsenatis* with a Km value of 0.3mM arsenate and Vmax of 7013 Umol/min [35]. Higher reduction efficiency of the isolate can be ascertained from the fact that it can actively reduce 88.2% of the initial 2mM arsenate [As(V)] to arsenite [As(III)] over a period of 72 hours. High activity of the enzyme leads to the conversion of arsenate to more mobile arsenite in the shallow aquifers that leads to its accumulation over a time period and could be one of the major reason for the increasing carcinogenic development in the northeastern region.

In a transitory, the production of siderophore by bacteria helps in the mobilization of the sedimentary arsenate by displacing iron from the iron-arseno compounds forming a soluble Fe^+3^ ion. Increase concentration of arsenate in surrounding milieu competes with the phosphate ion and structural homology of the arsenate with phosphate gives an added advantage to enter the cellular system through *pit/pst* phosphate transporter channel. Cellular arsenate is then converted to arsenite by arsenate reductase enzyme and soon effluxes out of the system through arsenite transporter channel to maintain the cellular homeostasis (**Fig. 8**) [19]. This essentially increases the concentration of both arsenite and arsenate in the aquifer system and eventually increases the arsenic contamination in the aquifers impacting the geocycle.

**Figure 8:**
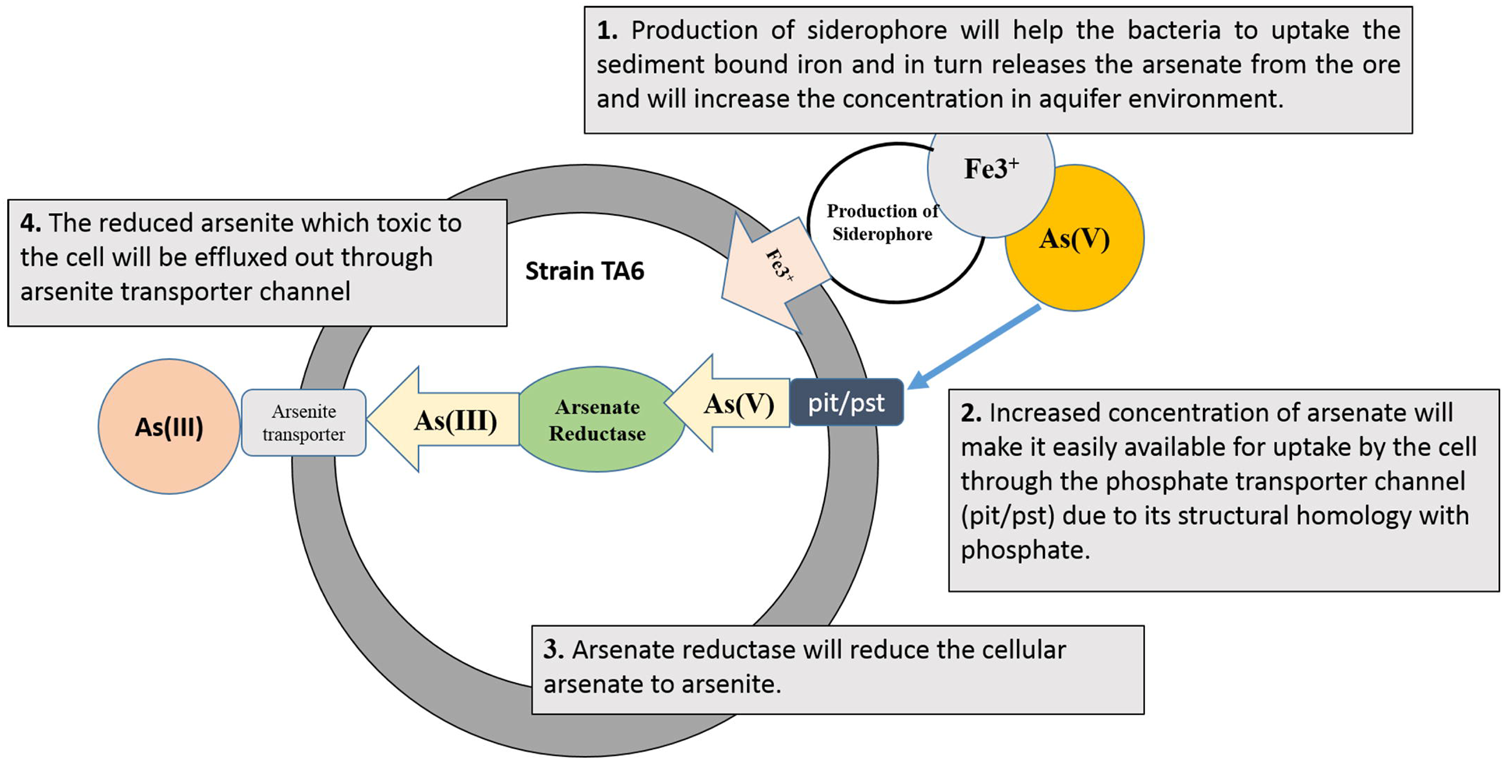
Graphical abstract representing the possible involvement of the strain in arsenic geocycle.

## Acknowledgment

Authors are grateful towards UGC for the Rajiv Gandhi National Fellowship grant to carry out the research work and all the faculty members of Department of Agricultural Biotechnology, Assam Agricultural University and Centre for Studies in Biotechnology, Dibrugarh University for their continuous support and guidance during the work.

## Reference

[1] Devi, N.L., Chandra, I., Shihua, Q., 2009. Recent Status of Arsenic Contamination in Groundwater of Northeastern India-A Review. Rep. Opin., 2009, 1, 22–32.

[2] Today, N., 2017. Water Sources in 23 Districts of Assam Contaminated with Arsenic and Fluoride □» Northeast Today 2017.

[3] Das, S., Bora, S., Lahan, J., Barooah, M., et al., 2014. GROUNDWATER ARSENIC CONTAMINATION IN NORTH EASTERN STATES OF INDIA. J. Environ. Res. Dev. J. Environ. Res. Dev., 2014, 9.

[4] Oremland, R.S., Stolz, J.F., 2005. Arsenic, microbes and contaminated aquifers. Trends Microbiol., 2005, 13, 45–49.

[5] Mukhopadhyay, R., Rosen, B.P., Phung, L.T., Silver, S., 2002. Microbial arsenic: From geocycles to genes and enzymes. FEMS Microbiol. Rev., 2002, 26, 311–325.

[6] Sarkar, A., Kazy, S.K., Sar, P., 2013. Characterization of arsenic resistant bacteria from arsenic rich groundwater of West Bengal, India. Ecotoxicology, 2013, 22, 363–376.

[7] Wang, G., Huang, Y., Li, J., 2011. [Bacteria live on arsenic analysis of microbial arsenic metabolism--a review]. Wei Sheng Wu Xue Bao, 2011, 51, 154–60.

[8] Lloyd, J.R., Oremland, R.S., 2006. Microbial transformations of arsenic in the environment: From soda lakes to aquifers. Elements, 2006, 2, 85–90.

[9] Kraemer, S.M., 2004. Iron oxide dissolution and solubility in the presence of siderophores. Aquat. Sci. - Res. Across Boundaries, 2004, 66, 3–18.

[10] Ghosh, P., Rathinasabapathi, B., Teplitski, M., Ma, L.Q., 2015. Bacterial ability in AsIII oxidation and AsV reduction: Relation to arsenic tolerance, P uptake, and siderophore production. Chemosphere, 2015, 138, 995–1000.

[11] Behari, J.R., Prakash, R., 2006. Determination of total arsenic content in water by atomic absorption spectroscopy (AAS) using vapour generation assembly (VGA). Chemosphere, 2006, 63, 17–21.

[12] Simeonova, D.D., LiÃ¨vremont, D., Lagarde, F., Muller, D.A.E., et al., 2004. Microplate screening assay for the detection of arsenite-oxidizing and arsenate-reducing bacteria. FEMS Microbiol. Lett., 2004, 237, 249–253.

[13] Aggett, J., Aspell, A.C., 1976. The determination of arsenic(III) and total arsenic by atomic-absorption spectroscopy. Analyst, 1976, 101, 341.

[14] Gladysheva, T.B., Oden, K.L., Rosen, B.P., 1994. Properties of the Arsenate Reductase of Plasmid R773. Biochemistry, 1994, 33, 7288–7293.

[15] Krieg, N.R., Krieg, R.N., 2015. Bergey’s Man. Syst. Archaea Bact., John Wiley & Sons, Ltd, Chichester, UK, pp. 1–14.

[16] Saitou, N., Nei, M., 1987. The neighbor-joining method: a new method for reconstructing phylogenetic trees. Mol. Biol. Evol., 1987, 4, 406–25.

[17] Erickson, K., 2010. The Jukes-Cantor Model of Molecular Evolution. PRIMUS, 2010, 20, 438–445.

[18] Schwyn, B., Neilands, J.B., 1987. Universal chemical assay for the detection and determination of siderophores. Anal. Biochem., 1987, 160, 47–56.

[19] Sarkar, A., Kazy, S.K., Sar, P., 2013. Characterization of arsenic resistant bacteria from arsenic rich groundwater of West Bengal, India. Ecotoxicology, 2013, 22, 363–376.

[20] Alexander, D.B., Zuberer, D.A., 1991. Use of chrome azurol S reagents to evaluate siderophore production by rhizosphere bacteria. Biol. Fertil. Soils, 1991, 12, 39–45.

[21] Chetia, M., Chatterjee, S., Banerjee, S., Nath, M.J., et al., 2011. Groundwater arsenic contamination in Brahmaputra river basin: a water quality assessment in Golaghat (Assam), India. Environ. Monit. Assess., 2011, 173, 371–385.

[22] Mahanta, C., Choudhury, R., Basu, S., Hemani, R., et al., 2015. Safe Sustain. Use Arsenic-Contaminated Aquifers Gangetic Plain, Springer International Publishing, Cham, pp. 57–64.

[23] Sarma, K.P., Patil, S.N., Kachate, N.R., Marathe, N.P., 2014. Groundwater arsenic contamination in the Brahmaputra and the Barak flood plains of Assam Hydrogeological studies from some parts of Malkapur area of Buldhana district, Maharashtra. Hydrol Curr. Res, 2014, 5.

[24] Mukherjee, A., Sengupta, M.K., Hossain, M.A., Ahamed, S., et al., 2006. Arsenic contamination in groundwater: A global perspective with emphasis on the Asian scenario. J. Heal. Popul. Nutr., 2006, 24, 142–163.

[25] Islam, F., Gault, G., Bootham, C., Polya, D., et al., 2004. Role of metal-reducing bacteria in arsenic release from Bengal delta sediments. Nature, 2004, 430, 68.

[26] Das, S., Bora, S.S., Yadav, R.N.S., Barooah, M., 2017. A metagenomic approach to decipher the indigenous microbial communities of arsenic contaminated groundwater of Assam. Genomics Data, 2017, 12, 89–96.

[27] Silver, S., Phung, L.T., 2005. Genes and enzymes involved in bacterial oxidation and reduction of inorganic arsenic. Appl. Environ. Microbiol., 2005, 71, 599–608.

[28] Jackson, C., Jackson, E., Dugas, S., Gamble, K., William, S., 2003. Microbial transformations of arsenite and arsenate in natural environments. Recent Res Dev Microbiol, 2003, 7, 103–118.

[29] Silver, S., Budd, K., Leahy, K.M., Shaw, W. V, et al., 1981. Inducible plasmid-determined resistance to arsenate, arsenite, and antimony (III) in escherichia coli and Staphylococcus aureus. J. Bacteriol., 1981, 146, 983–96.

[30] Ji, G., Silver, S., 1992. Regulation and expression of the arsenic resistance operon from Staphylococcus aureus plasmid pI258. J. Bacteriol., 1992, 174, 3684–94.

[31] Srivastava, S., Verma, P.C., Chaudhry, V., Singh, N., et al., 2013. Influence of inoculation of arsenic-resistant Staphylococcus arlettae on growth and arsenic uptake in Brassica juncea (L.) Czern. Var. R-46. J. Hazard. Mater., 2013, 262, 1039–1047.

[32] Hammer, N.D., Skaar, E.P., 2011. Molecular Mechanisms of Staphylococcus aureus Iron Acquisition. Annu. Rev. Microbiol., 2011, 65, 129–147.

[33] Basavraj, N., Deepak, V., 2011. Medical applications of siderophores. Eur. J. Gen. Med., 2011, 8, 229–235.

[34] Del Giudice, I., Limauro, D., Pedone, E., Bartolucci, S., Fiorentino, G., 2013. A novel arsenate reductase from the bacterium Thermus thermophilus HB27:Its role in arsenic detoxification. Biochim. Biophys. Acta - Proteins Proteomics, 2013, 1834, 2071–2079.

[35] Krafft, T., Macy, J., 1998. Purification and characterization of the respiratory arsenate reductase of Chrysiogenes arsenatis. FEBS J., 1998, 255, 647–653.

